# Temperature affects the host range of *Rhabdochlamydia porcellionis* and the infectivity of *Waddlia chondrophila* and *Chlamydia trachomatis* elementary bodies

**DOI:** 10.1101/2022.09.02.506449

**Authors:** Bastian Marquis, Silvia Ardissone, Gilbert Greub

**Author notes:** Address correspondence to Gilbert Greub.

## Abstract

The *Rhabdochlamydiaceae* family is a recent addition to the *Chlamydiales* order. Its members were discovered in cockroaches and woodlice but recent metagenomics surveys demonstrated the widespread distribution of this family in the environment. It was moreover estimated to be the largest family of the *Chlamydiales* order based on 16S rRNA encoding gene diversity. Unlike most chlamydia-like organisms, no *Rhabdochlamydiaceae* could be co-cultivated in amoebae and its host range remains largely unknown. Here, we tested the permissivity of various mammalian and arthropod cell lines to determine the host range of *Rhabdochlamydia porcellionis*, the only cultured representative of this family. While a growth could initially only be obtained in the Sf9 cell line, lowering the incubation temperature of the mammalian cells from 37 °C to 28 °C allowed *R. porcellionis* to grow in those cells. Furthermore, a 6 h exposure to 37 °C was sufficient to irreversibly block the replication of *R. porcellionis*, suggesting that this bacterium either lost or never acquired the ability to grow at 37 °C. We next sought to determine if temperature would also affect the infectivity of elementary bodies. Although we could not purify enough bacteria to reach a conclusive result for *R. porcellionis*, our experiment showed that the elementary bodies of *Chlamydia trachomatis* and *Waddlia chondrophila* lose their infectivity faster at 37 °C than at room temperature. Our results demonstrate that members of the *Chlamydiales* adapt to the temperature of their host organism and that this adaptation can in turn restrict their host range.

**Importance:** The *Rhabdochlamydiaceae* family is part of the *Chlamydiales*, a bacterial order that includes obligate intracellular bacteria sharing the same biphasic developmental cycle. This family have been shown to be highly prevalent in the environment, particularly in freshwater and soil and despite being estimated to be the largest family in the *Chlamydiales* order, is only poorly studied. Members of the *Rhabdochlamydiaceae* have been detected in various arthropods like ticks, spiders, cockroaches and woodlice, but the full host range of this family is currently unknown. In this study, we showed that *R. porcellionis*, the only cultured representative of the *Rhabdochlamydiaceae* family cannot grow at 37 °C and is quickly inactivated at this temperature. A similar temperature sensitivity was also observed for elementary bodies of chlamydial species adapted to mammals. Our work demonstrates that some chlamydiae adapt to the temperature of their reservoir, making a jump between species with different body temperatures unlikely.

## Introduction

The *Chlamydiales* order includes obligate intracellular bacteria that share the same biphasic developmental cycle composed of an extracellular infectious stage, the elementary body, and an intracellular replicative form, the reticulate body (1). While historically restricted to the human pathogens of the *Chlamydiaceae* family (2,3), this order has seen a rapid expansion during the past decades as new species were discovered in the environment (4–12). Estimations based on sequences diversity in the 16S rRNA encoding gene predict hundreds of unknown family level lineages (5). It is now evident that far from being restricted to mammals, chlamydiae are highly prevalent in the environment and successfully adapted to different ecological niches and host organisms (5,13–15).

Since they diverged from their common ancestor hundreds of millions of years ago (3,16), the different families of the *Chlamydiales* order specialized for specific hosts, often losing the ability to infect other organisms in the process. For instance, species of the *Chlamydiaceae* family, while highly adapted to vertebrates, are seemingly unable to replicate in amoebae (17–19). Conversely, members of the *Parachlamydiaceae* family grow efficiently in amoebae but poorly, if at all, in mammalian and insect cell lines, likely due to their inability to inhibit apoptosis (20–24). Some other families, like the *Simkaniaceae* and *Waddliaceae* families, seem to have conserved wider host ranges, as cultured representatives can grow in a wide variety of cell types (19,21,25–27). It is however unclear how the *in-vitro* host range of these bacteria translates *in-vivo*, as immortalized cell lines grown in axenic medium in the absence of immune system is far from the condition expected in most multicellular organisms. The host range of some *Chlamydiales* is thus well determined, while it remains unknown for others.

The picture is clearer for the *Rhabdochlamydiaceae* family, a recent addition to the *Chlamydiales* order. Members of this family were initially discovered in cockroaches (28) and woodlice (29) and later detected in ticks and spiders (30–34), while a distant relative was recently identified in the amoeba *Dictyostelium discoideum* (35). Interestingly, *Rhabdochlamydiaceae* were also detected in patients suffering of respiratory infections (36–38) or inflammatory skin disorders (39), suggesting a potential pathogenic role of these bacteria. Despite being hypothesized to be the most diverse clade of the *Chlamydiales* order (5), the *Rhabdochlamydiaceae* family is poorly studied and only one species, isolated from the rough woodlouse, could be cultured so far (40). Similarly to *Chlamydia trachomatis, Rhabdochlamydia porcellionis* was shown to inhibit apoptosis and could not be cultured in amoebae (40), hinting at a specialization for multicellular organisms. In line with this hypothesis, several arthropod cell lines were shown to sustain the growth of *R. porcellionis* (40). Unlike the *Chlamydiaceae, Rhabdochlamydiaceae* were not associated to animal hosts but to soil and freshwater environments (13) and while various arthropods were demonstrated to be a reservoir for *Rhabdochlamydiaceae*, their full host range is still unknown.

We thus aimed at better determining the host range of this bacterium. As *Rhabdochlamydiaceae* are suspected to be pathogenic towards human beings, we were interested in determining whether *R. porcellionis* could grow in mammalian cell lines. We also tested the permissivity of several arthropod cell lines.

## Methods

### Cell culture

*Spodoptera frugiperda* ovarian epithelial cells (Sf9, ATCC CRL-1711) were cultured at 28 °C in Grace insect medium (Gibco, Thermo Fisher Scientific, Waltham, USA) supplemented with 10% fetal calf serum (FCS). *Aedes albopictus* cells (C6/36, ATCC CRL-1660) were cultured at 28 °C in presence of 5% CO_2_ in DMEM (PAN-Biotech, Aidenbach, Germany) supplemented with 10% FCS. *Ixodes ricinus* (IRE/CTVM19) cell lines were maintained at 28 °C in Leibovitz L-15 medium (Gibco, Thermo Fisher Scientific, Waltham, USA) supplemented with 10% tryptose phosphate broth (Gibco, Thermo Fisher Scientific, Waltham, USA), 20% FCS and 1% L-glutamine (Sigma-Aldrich, Buchs, Switzerland), as described in (41). Human pneumocytes (A549, ATCC CCL-185), mouse fibroblasts (McCoy, ATCC CRL-1696) and human endometrial cells (Ishikawa, gift of Dr. G. Canny) were cultured in DMEM supplemented with 10% FCS and grown at 37 °C in presence of 5% CO_2_. *Acanthamoeba castellanii* strain (ATCC 30010) was cultured at 25 °C in peptone-yeast extract-glucose (PYG) medium.

### Bacterial strains

The *R. porcellionis* strain was acquired from the DMSZ collection (DSM 27522) and cultivated in Sf9 cells. The infected cells were passed once a week and non-infected cells were added approximately every four passages to compensate for host cell death due to the presence of the bacteria. *Waddlia chondrophila* strain WSU 86-1044 (ATCC VR-1470) was co-cultivated with *A. castellanii* in PYG broth at 32 °C. Suspensions of EBs were collected at 7 days post-infection, diluted 10 times and used to infect fresh *A. castellanii. Chlamydia trachomatis* (ATCC VR-902B) was cultivated in McCoy cells in DMEM supplemented with 10% FCS and 1 μg/ml cycloheximide.

### Infection procedure

Cells were seeded in a 24-wells plate (Corning) at a density of 10^5^ or 3×10^5^ cells per well 2 hours before infection. The infection procedures were performed as described in previous publications (26,40).

For *R. porcellionis*, suspensions of infected Sf9 were subjected to a freeze-thaw cycle to disrupt the cells, followed by a filtration through a 5 μm pore filter to remove the debris. Plated cells were then infected with the filtrate at an MOI ^~^0.1-1 and centrifuged for 15 minutes at 130 g at room temperature, followed by an incubation of 30 minutes at 28 °C. The medium was then replaced to remove non-internalized bacteria.

*W. chondrophila* EBs were collected from the supernatant of *A. castellani* at 5 days post-infection. The supernatant was then filtered through a 5 μm pore filter to remove cell debris. Plated cells were infected with the filtrate at an MOI ^~^0.1-1 and centrifuged at 1790 g for 10 minutes at room temperature. After 30 minutes of incubation at either 28 °C (for Sf9 cells) or 37 °C (for mammalian cells), the medium was replaced to remove non-internalized bacteria.

*C. trachomatis* EBs were collected from the supernatant of infected McCoy cells at 3 days post-infection. The supernatant was then filtered with a 5 μm pore filter. Plated cells were infected with the filtrate and centrifuged at 900 g for 15 minutes at room temperature. The infected cells were then incubated for 30 minutes at 37 °C and the medium was replaced to remove non-internalized bacteria.

After the infection, the cells were incubated at their usual growth temperature, unless specified otherwise. The samples were collected at various timepoints for quantification by qPCR and immunofluorescence staining.

### Infection cycle duration

Sf9 cells were plated in a 24-wells plate at a density of 10^5^ cells per well, infected with *R. porcellionis* at an MOI ^~^0.1-1 and incubated at 28 °C. The supernatant from the infected cells was collected at 0, 2, 4, 6, 8 and 10 days post infection and centrifuged at 300g for 5 minutes to pellet debris and detached cells. The supernatant containing EBs was then used to infect fresh Sf9 cells pre-plated in a 24-wells plate at a density of 10^5^ cells per well. Samples were taken at the end of the infection procedure and at 6 days post-infection. DNA was then extracted and the number of genome copies was measured by qPCR.

### Inclusion forming unit (IFU) quantification

Cells were plated in a 24-wells plate at a density of 3×10^5^ cells per well two hours before the infection and were then infected with serial tenfold dilutions of EBs suspensions. After the initial 30 minutes of incubation, the old medium was replaced with fresh medium supplemented with 1 μg/ml of cycloheximide (42). The cells infected with *W. chondrophila* or *C. trachomatis* were incubated for 24 hours at 37 °C with 5% CO_2_, while the cells infected with *R. porcellionis* were incubated for 6 days at 28 °C, with 5% CO_2_ for mammalian cells. The cells were then prepared for immunofluorescence and the proportion of infected cells was determined using an epifluorescence microscope. At least 100 cells were counted for each condition.

### Effect of temperature on EBs infectivity

Elementary bodies were collected following the same procedure used for infections. The filtrate was diluted 1:1 in PYG for *W. chondrophila*, 1:1 in DMEM supplemented with 10% FCS for *C. trachomatis* and 1:1 in Grace medium supplemented with 10% FCS for *R. porcellionis*. The dilutions of elementary bodies were then incubated at 20 °C or 37 °C in 24-well plates. In the case of *C. trachomatis*, the plate was incubated in presence of 5% CO_2_. IFUs were quantified using McCoy cells immediately after the dilution in fresh medium or after 1, 2, 3 or 4 days of incubation.

### Effect of incubation temperature on growth

Sf9 cells were plated in a 24-well plate at a density of 10^5^ cells per well, infected with *R. porcellionis* at an MOI of ^~^0.1-1 and incubated for 48 hours at 28 °C. The plates were then incubated at 37 °C for 6, 12, 24 or 48h before being switched back to 28 °C for four additional days. Samples were taken for bacterial growth quantification by qPCR right after the switch to 28 °C and after the subsequent 4 days of incubation.

### Quantitative PCR

Genomic DNA was extracted using the Wizard SV Genomic DNA Purification Kit (Promega, Dübendorf, Switzerland) following the manufacturer protocol. Quantitative PCR for *R. porcellionis* (36) or *W. chondrophila* (43) was performed on 5μL of genomic DNA with iTaq Supermix (Bio-Rad, Cressier, Switzerland), 200 nM primers (WadF4 5′-GGCCCTTGGGTCGTAAAGTTCT-3′ and WadR4 5′-CGGAGTTAGCCGGTGCTTCT-3′ for *W. chondrophila*; RcF 5′GACGCTGCGTGAGTGATGA-3′ and RcR 5′-CCGGTGCTTCTTTACGCAGTA-3′ for *R. porcellionis*) and 100 nM probe (WadS2 5′-FAM-CATGGGAACAAGAGAAGGATG-BHQ1-3′ and RcS 5′-FAM-CTTTCGGGTTGTAAAACTCTTTCGCGCA-BHQ1-3′). The cycling conditions were identical for both qPCRs: 3 min at 95 °C, 40 cycles of 15 seconds at 95 °C and 1min at 60 °C. The qPCRs were performed on a QuantStudio3 real-time PCR system (Applied Biosystems, Thermo Fisher Scientific, Waltham, USA).

### Immunofluorescence staining

Infected cells grown on glass coverslips were fixed with ice-cold methanol for 5 minutes at different timepoints after the infection. Cells were then washed three times with PBS and incubated for at least two hours in PBS with 0.1% saponin, 0.04% NaN3 and 10% FCS (blocking solution). The coverslips were then incubated at room temperature for 2 hours in blocking solution with rabbit *anti-Simkania negevensis* antibodies (25) (dilution at 1:1000), rabbit anti-*Waddlia chondrophila* antibodies (44) (dilution 1:1000) or goat antibodies targeting the major outer membrane protein of *Chlamydia trachomatis* (dilution 1:1000) (LSBio, Seattle, USA). Anti-*Simkania* antibodies were used to detect *R. porcellionis*, as antibodies raised against a chlamydial species often cross-react with related species (40,45). After the incubation with the primary antibody, the coverslips were washed three times in PBS with 0.1% saponin and incubated for one hour at room temperature in blocking solution with 1.6 μg/ml DAPI dilactate (Molecular Probes, Thermo Fisher Scientific, Waltham, USA), 100 μg/ml Concanavalin A-Texas red conjugate (Invitrogen, Thermo Fisher Scientific, Waltham, USA) and Alexa-488 conjugated chicken anti-goat or goat anti-rabbit antibodies (1:1000 dilution) (Life Technologies, Thermo Fisher Scientific, Waltham, USA). The coverslips were then embedded in Moewiol (Sigma-Aldrich, Buchs, Switzerland) and kept in the dark at 4 °C until further use. The coverslips were examined with a confocal microscope Zeiss LSM 900 (Zeiss, Oberkochen, Germany).

### Statistical analysis

The results of this study are given as means with their standard deviation. The linear regression models with random effect were fitted using the lme4 package (46). All statistics were performed with R (v4.2.0).

## Results

### *R. porcellionis* has a limited host range and a long replication cycle

To study the host range of *R. porcellionis* in mammalian cells, we tested the permissivity of pneumocytes (A549), endometrial cells (Ishikawa) and fibroblasts (McCoy). The first two cell lines are indeed known to be permissive to different *Chlamydia*-like organisms (25,27), while McCoy cells are frequently used to propagate members of the *Chlamydiaceae* family. In addition to the Sf9 cell line, already used for the subculture of *R. porcellionis* (40), we tested a tick cell line (IRE/CTVM19) and a mosquito cell line (C6/36) as those were derived from organisms likely closer to the natural reservoir of the *Rhabdochlamydiaceae* family than mammalian cell lines.

Growth could be observed in Sf9 (Fig 1A), with a doubling time of 20.4h (sd=0.4h), comparable to that of *Simkania negevensis* (21h) but longer than *Waddlia chondrophila* (4h) in the same cell line (21,25). Immunofluorescence showed heavily infected Sf9 cells at 6 days post-infection (Fig 1C), however, reticulate bodies are disseminated in the cytoplasm of the host cell and do not seem to be enclosed in an inclusion (Fig. 1C, Fig. 4C, Supp. Movie. 1). Interestingly, unlike *S. negevensis, R. porcellionis* could not grow in mammalian cells and, more surprisingly, failed to grow in any of the other arthropod cell lines (Fig 1A). While bacteria were internalized in those cells (Fig 1C), they either failed to initiate replication or formed aberrant bodies measuring more than 5 μm (in C6/36 cells).

**Figure 1.**
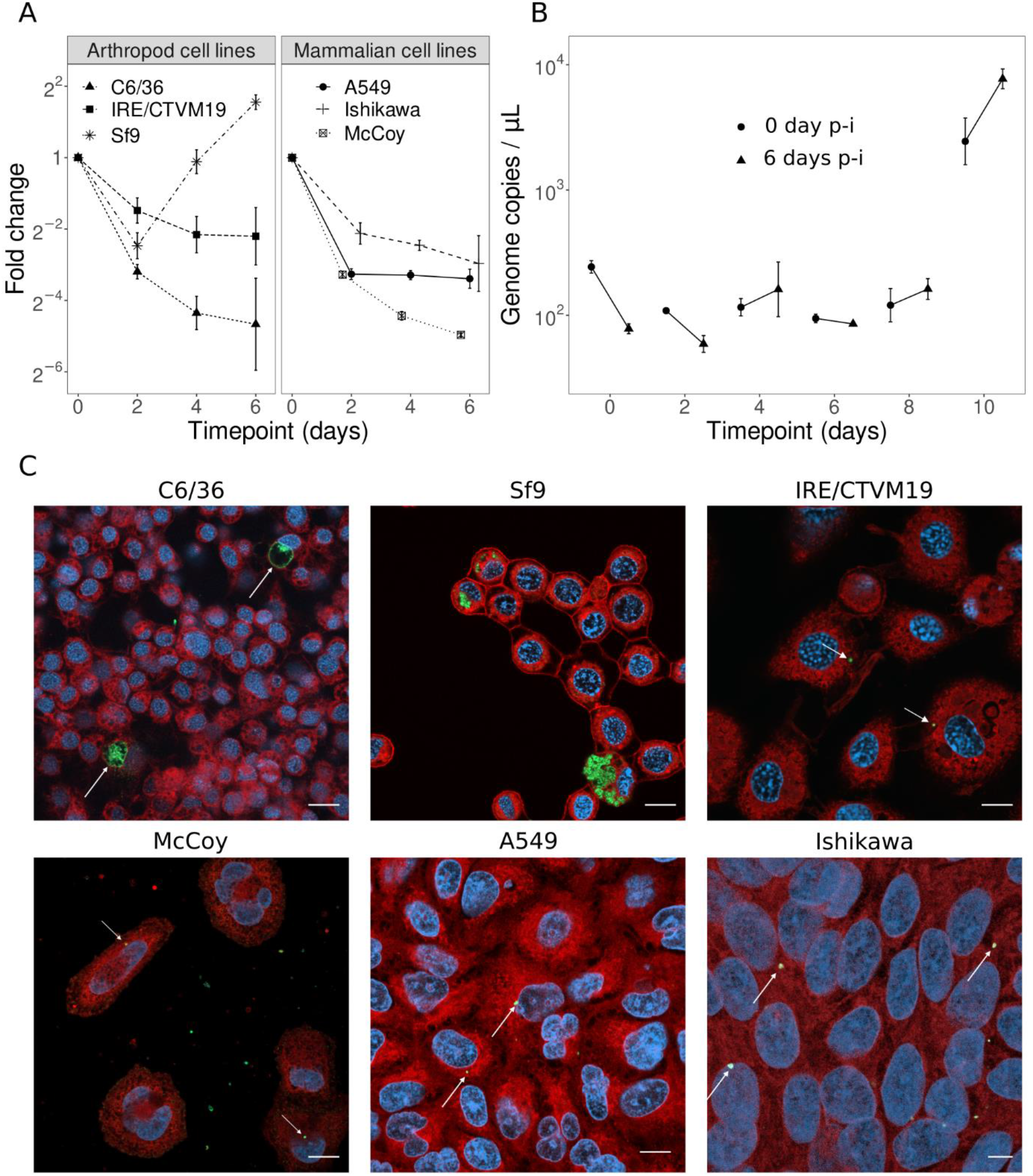
**(A)** Permissivity of arthropod and mammalian cell lines to *R. porcellionis*. The y-axis represents the log2 fold change of the number of genome copies per μL compared to the initial timepoint (mean +/− standard deviation, n=2) **(B)** Infectious progeny production. The supernatant from Sf9 cells infected with *R. porcellionis* was collected every other day (time points indicated on the x axis) and used to infect fresh Sf9 cells. The cells were then collected at zero (dots) and 6 days (triangles) post infection to quantify bacterial growth by qPCR (mean +/− standard deviation, n=2) **(C)** *R. porcellionis* in mammalian and arthropod cell lines at 6 days post-infection. Growth could only be seen in Sf9 cells. The reticulate bodies seem to replicate directly in the cytoplasm, without being enclosed in an inclusion. The enlarged bodies in the C6/36 cell lines are likely aberrant bodies. Bacteria appear to have been internalized in all the other cell lines, but failed to replicate. White arrows indicate enlarged bacteria in C6/36 cells and internalized EBs in the other cell lines. Cells were stained with concanavalin A (red), DAPI (blue) and anti-*Simkania* antibody (green), scale bar: 10 μm.

**Figure 2.**
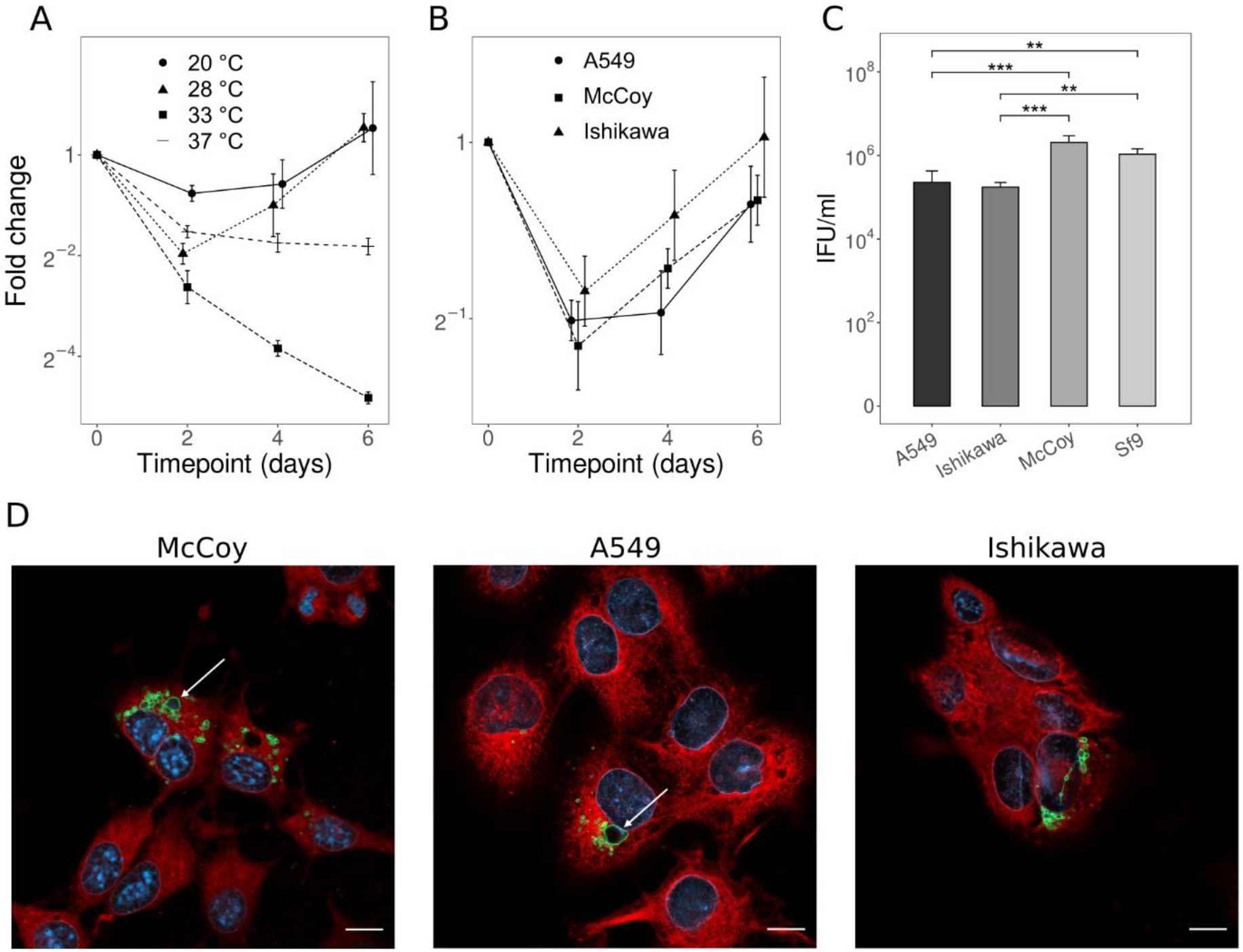
*R. porcellionis* grows in mammalian cells incubated at 28 °C **(A)** Growth kinetics of *R. porcellionis* in Sf9 incubated at 20, 28, 33 and 37 °C. **(B)** Growth kinetics of *R. porcellionis* in mammalian cells, incubated at 28 °C. In both **(A)** and **(B)**, the y-axis represent the log2 fold change compared to time 0 (mean +/− standard deviation, n=3) **(C)** IFU count in different cell lines infected with the same suspension of *R. porcellionis* (mean +/− standard deviation, n=3). All cells were all grown at 28 °C. A one-way ANOVA revealed that there was a statistically significant difference between at least two cell lines (p-value=0.0002). The plot also shows the results of Tukey HSD test for the pairwise comparison of the different cell lines (** < 0.01, *** < 0.001). **(D)** McCoy, A549 and Ishikawa cells infected with *R. porcellionis* and incubated at 28 °C. The two enlarged bodies in McCoy and A549 cells are likely aberrant bodies (white arrow). Cells were fixed 6 days post infection and stained with concanavalin A (red), DAPI (blue) and anti-*Simkania* antibodies (green). Scale bar: 10 μm.

**Figure 3.**
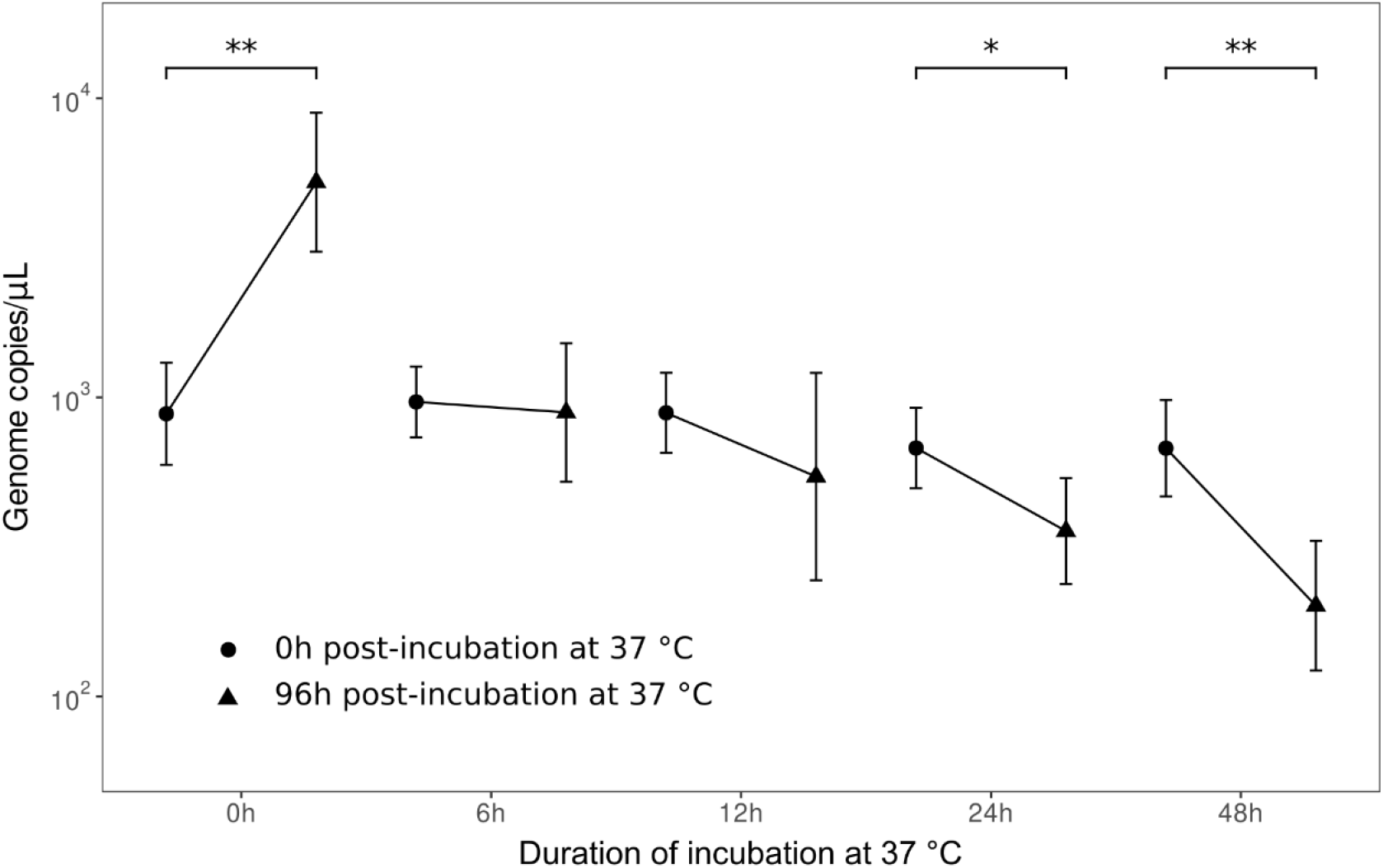
Effect of a transient exposure to 37 °C on the replication of *R. porcellionis*. The graph shows the number of bacteria (genome copies/μl, as determined by qPCR) immediately after the exposure to 37°C (dots) and after 4 days of recovery at 28°C (triangles). The plot also shows for which timepoints the log-ratio of the genome copies at the 96h and 0h was significantly different from 0. The results were corrected for multiple testing with the Holm step-down procedure (mean +/− standard deviation, n=3, * < 0.05, ** < 0.001, t-test).

**Figure 4.**
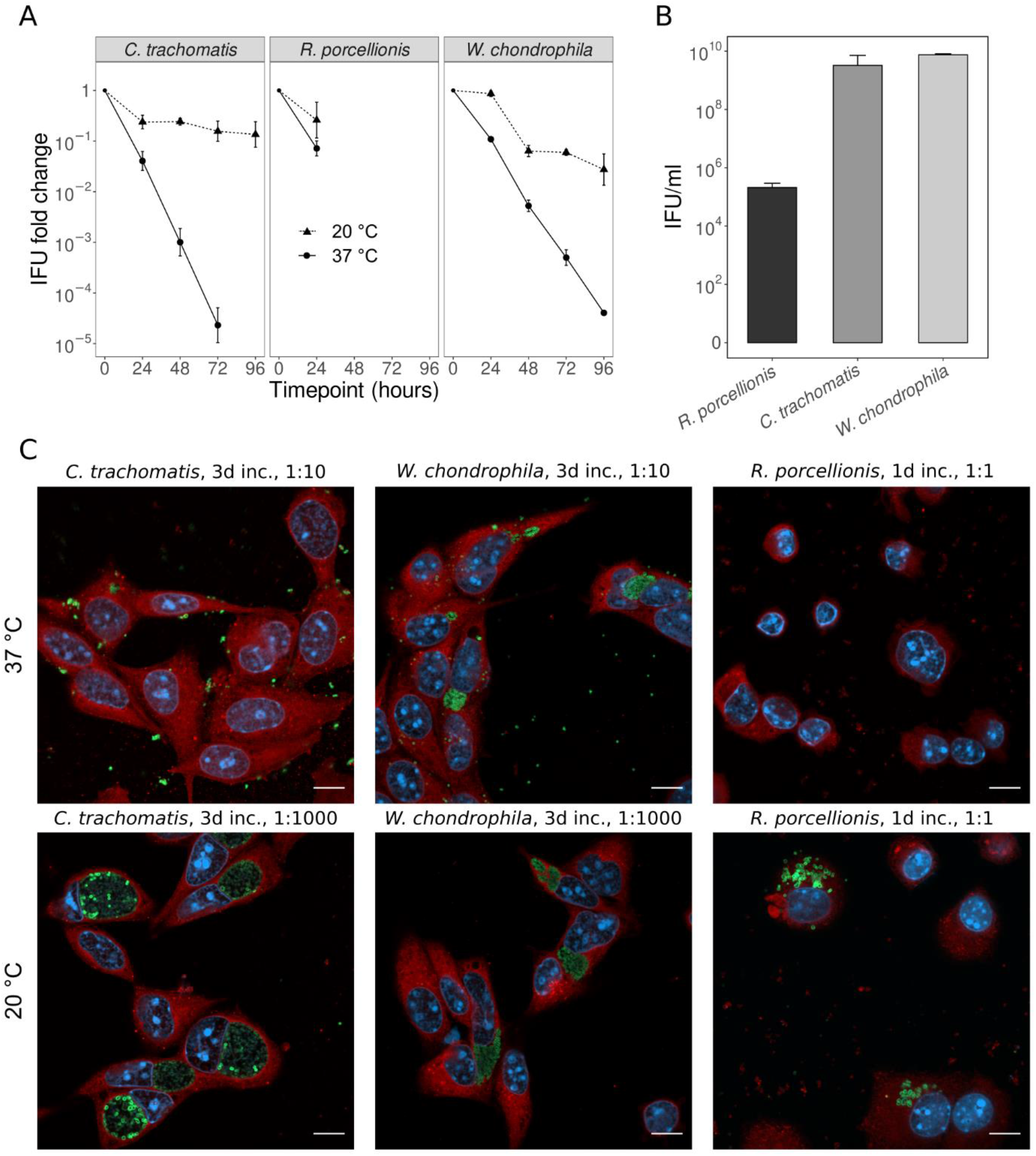
Effect of different incubation temperatures on the infectivity of elementary bodies. **(A)** Evolution of the number of IFU after incubation at 20 °C or 37 °C, normalized to the initial IFU count (mean +/− standard deviation, n=3) **(B)** IFU counts at the initial timepoint (mean +/− standard deviation, n=3) **(C)** Confocal images of McCoy cells infected with serial dilutions of EBs incubated at 20 °C or 37 °C. This highlights the deleterious effect of an incubation at 37 °C for the EBs of all three species. Interestingly, the inclusions formed by EBs incubated at 37 °C also tended to be smaller. The difference between the well-formed inclusions for *C. trachomatis* and *W. chondrophila* and the dissemination of RBs in the host cytoplasm for *R. porcellionis* is also striking. Cells were fixed at 1 (for *C. trachomatis* and *W. chondrophila)* or 6 days (for *R. porcellionis*) post-infection and stained with Concanavalin A (red), DAPI (blue) and antibodies against the different bacteria (green). Scale bar: 10μm. Inc.: incubation duration.

In line with the long doubling time, the release of infectious progeny from infected cells also occurs late after the initial infection. A significant increase of infectious particles in the supernatant of infected cells was indeed observed only 10 days post infection (Fig 1B), whereas in the case *C. trachomatis, W. chondrophila* or *S. negevensis* the infectious cycle is completed within 2-5 day*s*.

### *R. porcellionis* is unable to replicate at 37 °C

*Rhabdochlamydiaceae* have been detected in various arthropods such as ticks (47,48), spiders (49), cockroaches (28) and woodlice (29). As those organisms are poikilothermic and have a lower body temperature than mammals, we reasoned that the *Rhabdochlamydiaceae* family might have adapted to the lower temperature of its host organisms and have either lost or never acquired the ability to grow at 37 °C. To test this hypothesis, we infected Sf9 cells with *R. porcellionis* and incubated them at 20, 28, 33 and 37 °C. As a control for the effect of the temperature on host cells, we infected Sf9 cells with *W. chondrophila*, which grows well at 37°C in different cell lines (21,26,27), and incubated them at 28 and 37 °C. *R. porcellionis* grew at 20 and 28 °C, but not at 33 or 37 °C (Fig 2A). Conversely, *W. chondrophila* replicated more efficiently at 37 °C than at 28 °C (Supplementary Fig 1). This suggests that a loss of permissivity of Sf9 cells at 37°C is unlikely, and that the effect of temperature observed for *R. porcellionis* was rather due to a loss of infectivity and/or replication of the bacteria.

To further test this hypothesis, we infected A549, Ishikawa and McCoy cells with *R. porcellionis* and lowered the incubation temperature to 28 °C. This allowed the growth of *R. porcellionis* (Fig 2B) in mammalian cells, although the doubling time was significantly longer in Ishikawa and McCoy cells than in Sf9 cells (Table 1). Aberrant bodies were also frequently observed in mammalian cells (Fig 2D). Altogether, this data confirms that temperature is a limiting factor for the host range of *R. porcellionis*, while the long doubling times and the presence of aberrant bodies suggest that the physiology of mammalian cells might be too different from the natural host of this bacterium to allow an efficient grow.

**Table 1:**
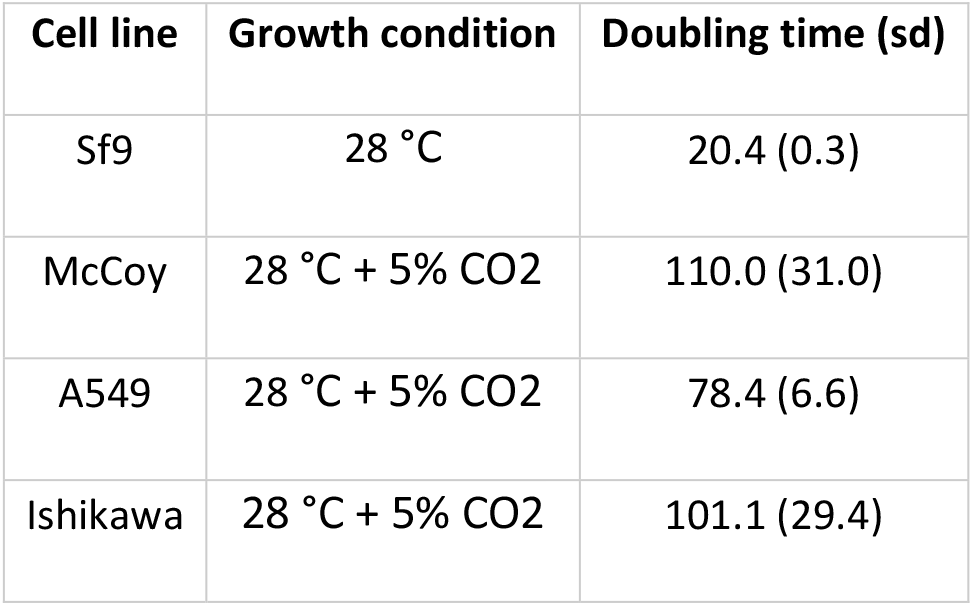
Doubling times of *R. porcellionis* in different cell lines. The doubling time in Sf9 cells differed significantly from the doubling time in McCoy (p=0.023) and Ishikawa (p=0.032) cells.

| Cell line | Growth condition | Doubling time (sd) |
| --- | --- | --- |
| Sf9 | 28 °C | 20.4 (0.3) |
| McCoy | 28 °C + 5% CO<sub>2</sub> | 110.0 (31.0) |
| A549 | 28 °C + 5% CO<sub>2</sub> | 78.4 (6.6) |
| Ishikawa | 28 °C + 5% CO<sub>2</sub> | 101.1 (29.4) |

Finally, we compared the infection efficiency of *R. porcellionis* by simultaneously infecting A549, McCoy, Ishikawa and Sf9 cells with the same dilutions of elementary bodies and measuring the IFU count at six days post-infection. As shown in Fig. 2C, the IFU count was significantly lower in Ishikawa and A549 compared to Sf9 cells. Surprisingly, there was no difference between McCoy and Sf9 cells, which might indicate a tropism of *R. porcellionis* for fibroblasts. The IFU count in Sf9 might however be under-estimated, as infected cells tend to detach from the coverslip.

### Transient exposure to 37 °C irremediably blocks the replication of *R. porcellionis*

Members of the *Chlamydiaceae* are known to enter a third non-replicative stage, the aberrant body, when exposed to stresses such as antibiotic exposure (50), nutrient deprivation (51) or heat shock (52). Aberrant bodies are typically described as non-replicating enlarged cells able to resume their regular cycle once the stress disappears, although their morphology was shown to vary in function of the stresses (53,54). We thus wondered whether *R. porcellionis* could similarly recover and resume its growth cycle after an exposure to 37 °C. To determine this, Sf9 cells at 2 days post-infection were incubated at 37 °C for various durations. The infected cells were then further incubated at 28 °C for four additional days to check for bacterial growth after stress removal. The growth was assessed by measuring the number of genome copies at the end of the incubation at 37 °C and after 4 days of recovery at 28 °C.

Our experiment shows that a transient exposure to 37 °C as short as 6h irreversibly blocks the replication of *R. porcellionis* (Fig. 3). The effect of the temperature shift seems to be more deleterious for longer incubation at 37 °C, as expected. However, the complete experiment lasted from 6 days, for the samples subjected to the 6h shift at 37°C, to 8 days, for the samples subjected to the 48h shift at 37°C. This difference of duration could also explain the more pronounced effect of longer incubations. Indeed, the number of genome copies at the end of the shift to 37°C was not affected by the duration of the exposure (p-value=0.126, one-way ANOVA), showing that it takes at least 48h for temperature to affect the number of genome copies. It is therefore possible that the apparent lack of difference between 96h and 0h for the 6 and 12h timepoints is due to the experiment not being long enough for the effect of temperature to manifest in terms of genome copies. The delay might be due to DNA being slowly released – and degraded by cytosolic DNases – from inactivated bacteria.

### Temperature affects the infectivity of elementary bodies

Given the effect of temperature on the replication of *R. porcellionis*, we wondered whether it would also affect the infectivity of elementary bodies. To determine this, we incubated elementary bodies of *R. porcellionis, W. chondrophila* and *C. trachomatis* at 20 °C (room temperature) or 37 °C and measured the number of IFU every day for four days. To be closer to natural conditions and have a similar EB purification method for all three species, we did not perform the freeze and thaw cycle and directly filtered the supernatant from infected Sf9 cells. In addition, as Sf9 cells tend to detach after several days of infection, we used McCoy cells grown at 28 °C for IFU quantification of *R. porcellionis* to avoid any bias due to cell detachment. We used a random intercepts mixed-effects linear model to predict the log-transformed count of IFUs based on an interaction variable between the duration of incubation in days and the temperature. When none of the counted cells was infected, we conservatively assumed an IFU count corresponding to one infected cell in 100.

As shown in Fig. 4A, *R. porcellionis* EBs incubated for 24h at 37 °C were less infectious than their counterpart incubated at 20 °C, although the incubation temperature did not significantly predict the IFU count (R^2^=0.77, beta for the interaction term = −0.55, p-value = 0.06). The interaction term means that for every day of incubation at 37 °C, the IFU count will be reduced by a factor of 3.55 (10^0.55^) compared to the same incubation at room temperature. Due to the low initial quantity of infectious particles (Fig. 4B), no infected cell could be observed after the first timepoint. In the absence of any practical alternative to compensate for the 10’000× difference of initial infectious particles count with *C. trachomatis* and *W. chondrophila* (Fig. 4B), we did not further repeat the experiment with *R. porcellionis*.

Surprisingly, the trend observed for *R. porcellionis* was confirmed for *C. trachomatis* and *W. chondrophila*. As both bacteria thrive at 37 °C, we did not expect their elementary bodies to be affected by temperature. This proved to be false, as the temperature of incubation significantly affected the IFU count for both *W. chondrophila* (R^2^=0.97, beta = −0.68, p-value = 3.27×10^-11^) and *C. trachomatis* (R^2^=0.94, beta = −1.06, p-value = 2.08×10^-10^). Compared to an incubation at room temperature, the IFU count of *C. trachomatis* and *W. chondrophila* is thus predicted to be reduced by a factor of 11.48 and 4.79 for every day of incubation at 37 °C, respectively.

## Discussion

Several factors are known to influence the host range of members of the *Chlamydiales* order. The ability to inhibit apoptosis has for instance been suggested to be a hallmark of the chlamydiae infecting mammals (20,23), while organisms adapted to mammals seem to have lost the ability to grow in amoebae, for some unknown reason (3,19). Adaptation for specific temperature ranges has already been demonstrated to be an important determinant for the host range of the *Parachlamydiaceae* family (22,55,56). In the present study, we demonstrated that this impact of temperature extends to other families of the *Chlamydiales* order by showing that *R. porcellionis* likely specialized for the temperature ranges encountered in its arthropod hosts. As a consequence of this specialization, *R. porcellionis* is quickly inactivated if exposed to the body temperature of mammals, making it unlikely for this bacterium to be pathogenic for humans and restricting its host ranges to organisms with a temperature lower than 33 °C. Our results are consistent with previous evidences and suggest that *Chlamydiales* do co-evolve with their host and adapt to their body temperature (3,55,56). This adaptation might in turn become a limiting factor for the host range, as shown for *R. porcellionis* and *P. acanthamoebae* that were only able to grow in mammalian cells if those were incubated at lower temperatures. Mammals are thus unlikely to serve as a reservoir for either *R. porcellionis* or *P. acanthamoebae*.

*R. porcellionis* seems to have a tropism for specific cell types. In this study, it did not multiply in mosquito (C6/36) or tick cell lines (IRE/CTVM19), contrarily to a previous study where a growth could be obtained in a fruit fly cell line (S2) and in another mosquito cell line (Aa23-T) (40). However, those cell lines were derived from embryos and larvae and their exact cell type is unknown. It is therefore possible that this discrepancy is due to the C6/36 and Aa23-T cell lines originating from different tissues, some of which may not be permissive. This hypothesis is coherent with the apparent limitation of *R. porcellionis* distribution to the hindgut epithelium of woodlice (57). The discrepancy in the permissivity of the different arthropod cell lines contrasts with the consistent growth in the 3 mammalians cell lines at 28 °C. While the long doubling times and frequent aberrant bodies observed in those cells suggest that they do not offer ideal conditions for the growth of *R. porcellionis*, the ability of this bacterium to grow in mammalian cells indicates that it does not rely on arthropod-specific receptors for its adherence and internalization. Both the Ishikawa and the A549 cells were isolated from carcinoma, while McCoy cells are mouse fibroblasts. As Sf9 also exhibit an epithelial morphology, it is possible that *Rhabdochlamydia* has a tropism for epithelial cells and fibroblasts.

The temperature sensitivity of the elementary bodies of *C. trachomatis* and *W. chondrophila* is a surprising finding. A similar sensitivity is likely for *R. porcellionis*, but due to the difficulty of purifying enough bacteria, we were unable to confirm it. Although several studies already hinted at the effect of temperature on the infectivity in *Parachlamydia acanthamoebae* (58) and *Chlamydia pneumoniae* (58), such a marked effect was surprising in *Chlamydia trachomatis*. This bacterium is indeed a frequent human sexually transmitted pathogen that should be well-adapted to the temperature of mammals. The situation is less clear for *W. chondrophila*. Its natural reservoir is unknown, so it cannot be excluded that this bacterium usually thrives in ecological niches with lower temperature. But its isolation from a bovine abortus, its ability to grow at 37 °C and its association with miscarriages (59) suggest that, like *C. trachomatis*, it should be adapted to mammalian body temperatures. Assuming that the temperature sensitivity is a constitutive trait of elementary bodies, it may be that chlamydiae infecting warm blooded animals had to develop faster infectious cycles to prevent newly differentiated elementary bodies from being exposed to high temperatures for long durations while still in the inclusion. Elementary bodies are indeed produced asynchronously and can be isolated as early as 24h post-infection in *C. trachomatis*. The induction of a lytic infection when *Parachlamydia* is exposed to temperatures higher than 30 °C (60) could be similarly explained. The bacterium might induce the lysis of its host cell to accelerate the release of elementary bodies. The infectious cycle duration would therefore be directly affected by the temperature of the host.

*C. trachomatis* has been shown to require either glucose-6-phosphate or ATP to maintain its infectivity in extracellular medium, as it cannot metabolize glucose (42,61). This is consistent with the loss of infectivity we observed at 37 °C, as none of those molecules is present in the incubation medium we used, but fails to explain why the infectivity of *C. trachomatis* is preserved at room temperature. Moreover, unlike *C. trachomatis, W. chondrophila* has a complete glycolytic pathway (62). Its infectivity should therefore be preserved at both temperatures. In line with the hypothesis that metabolism is a key determinant of elementary body infectivity (42,61), it is possible that elementary bodies only marginally rely on external supplies of nutrients and use their own store of glycogen. Glycogen particles were indeed observed in elementary bodies and enzymes necessary for its synthesis are well conserved in the *Chlamydiales* order (63). Incubating elementary bodies at higher temperatures could therefore induce a faster metabolic rate and deplete the glycogen stores of elementary bodies. Alternatively, temperature has also been demonstrated to influence the activity of the type 3 secretion system (64). In particular, incubation at 37 °C was shown to induce the secretion of the Tarp effector protein compared to the same incubation at 4 °C, in the presence of liposomes. Depletion of effectors could therefore lead to the observed loss of infectivity. While in the same work (64), the activation of the type 3 secretion system did not seem to affect elementary bodies infectivity, the experiments only lasted 30 minutes. A loss of infectivity due to the activation of the type 3 secretion system therefore cannot be excluded for longer incubation durations. Our medium did not contain liposomes, but is also possible that FCS or cell debris smaller than 5 μm could act in concert with the temperature to induce the activation of the type 3 secretion system. Additional experiments are thus needed to determine the exact mechanism of this observed loss of infectivity.

In conclusion, our work show that as chlamydiae evolved from protists symbionts and adapted to new hosts, they also had to adapt to new ranges of temperatures. This specialization for specific temperatures in turn precludes species jump between organisms whose body temperatures are too different. Interestingly, elementary bodies apparently escaped this adaptation and kept similar temperature sensitivities, which may explain the fastest cycles observed in chlamydiae infecting organisms with high body temperatures. Finally, our results also suggest that various incubation temperatures should be tried when attempting to isolate chlamydiae from environmental samples.

## Acknowledgments

Bastian Marquis is funded by the Jürg Tschopp MD-PhD scholarship

## Contributions

Conceptualization: BM, GG

Data curation: BM

Formal Analysis: BM

Funding acquisition: BM, GG

Investigation: BM, GG, SA

Methodology: BM, SA

Project administration: GG

Resources: GG, SA

Software: N/A

Supervision: GG, SA

Validation: GG, SA

Visualization: BM

Writing – original draft: BM

Writing – review & editing: GG, SA

